## Supplementary Material for "The juxtamembrane linker of synaptotagmin 1 regulates Ca^2+^ binding via liquid-liquid phase separation"

*Corresponding author: Edwin R. Chapman

This PDF file includes:

Supplementary Figures 1-14

Supplementary Tables 1-4

**Supplementary Figures**

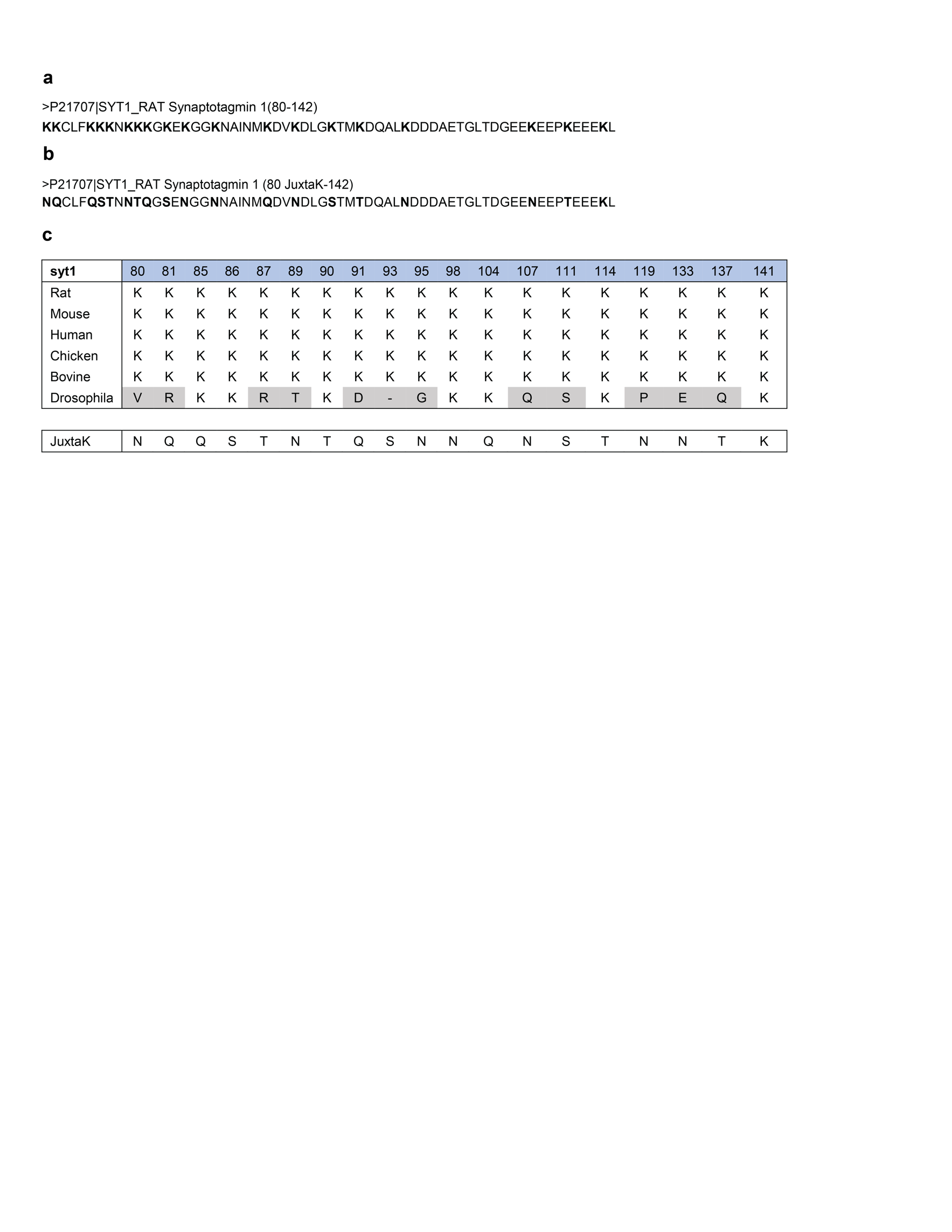

**Supplementary Fig. 1.** **Conservation of lysine residues in the syt1 juxtamembrane linker**. **a, b** Sequences of the juxtamembrane linkers (residues 80-142) from WT (*Rattus norvegicus*) and JuxtaK mutant, in which the lysine residues have been substituted with polar residues. **c** Alignment of all lysine residues in the juxtamembrane linkers of syt1 from multiple species, revealing strong conservation, except in flies. At the bottom, the amino acids used to substitute all but one lysine residue are shown. Aligned using ClustalOmega^1^.

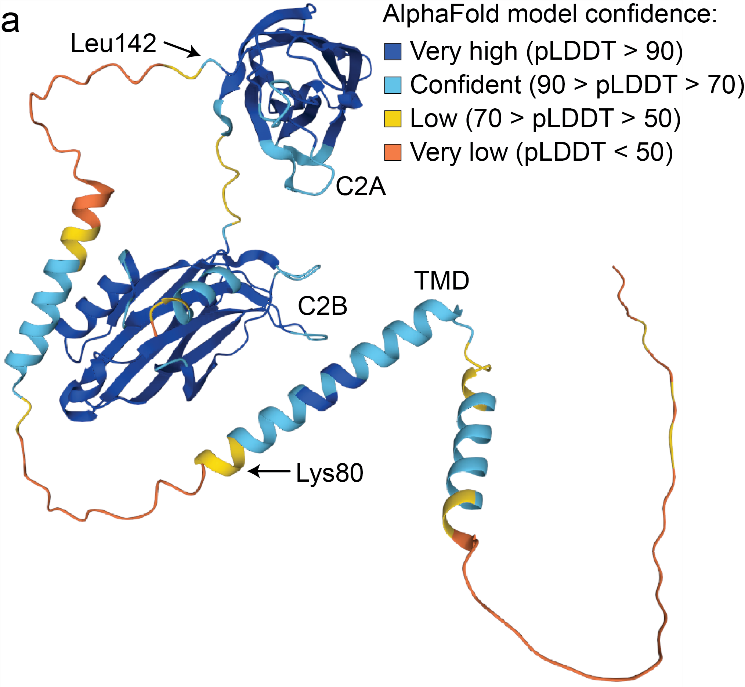

**Supplementary Fig. 2. Color-coded AlphaFold predicted syt1 structure, along with pLDDT scores, illustrating both folded and unstructured domains.**

**a** AlphaFold^2,3^ predicts that the juxtamembrane linker segment, residues Lys80 to Leu142, is mostly unstructured with a pLDDT score less than 50. There is a short sequence in the juxtamembrane linker segment that is predicted to be helical, with a higher pLDDT score. pLDDT: predicted local distance difference test per-residue confidence score.

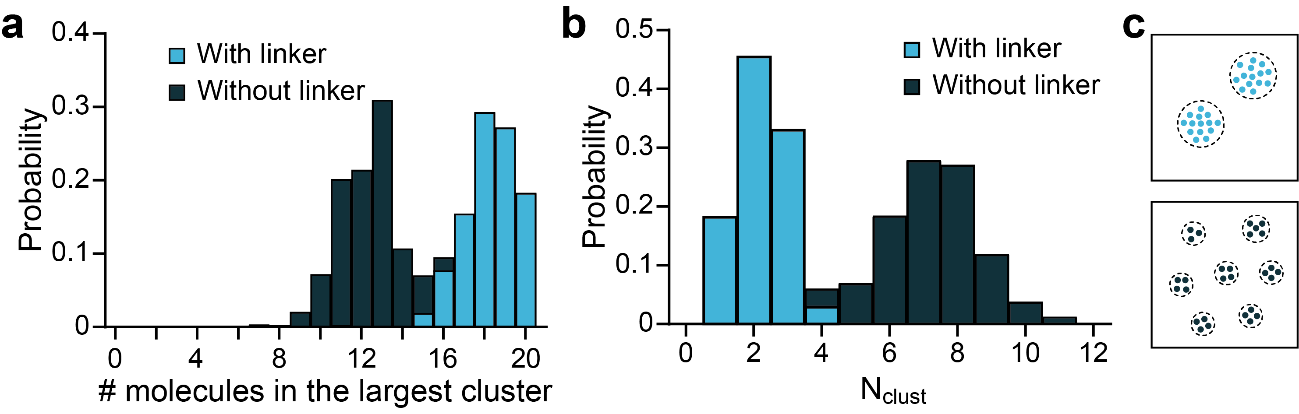

**Supplementary Fig. 3. The syt1 juxtamembrane linker drives LLPS.**

**a** Probability distribution of the number of syt1 molecules, with and without the juxtamembrane linker, in the largest cluster observed along the MD trajectories. Syt1 molecules with the juxtamembrane linker tend to form larger clusters, indicating that the presence of linker drives LLPS; n = 2 trajectories. **b** Probability distribution of the number of syt1 clusters (N_clust_), with and without the juxtamembrane linker. Removal of the linker results in the formation of a larger number of smaller clusters; n = 2 trajectories. **c** Illustration of syt1 clusters; a small number of large clusters form with an intact juxtamembrane linker (upper); a larger number of smaller clusters form in the absence of the linker (lower). Source data are provided as a Source Data file.

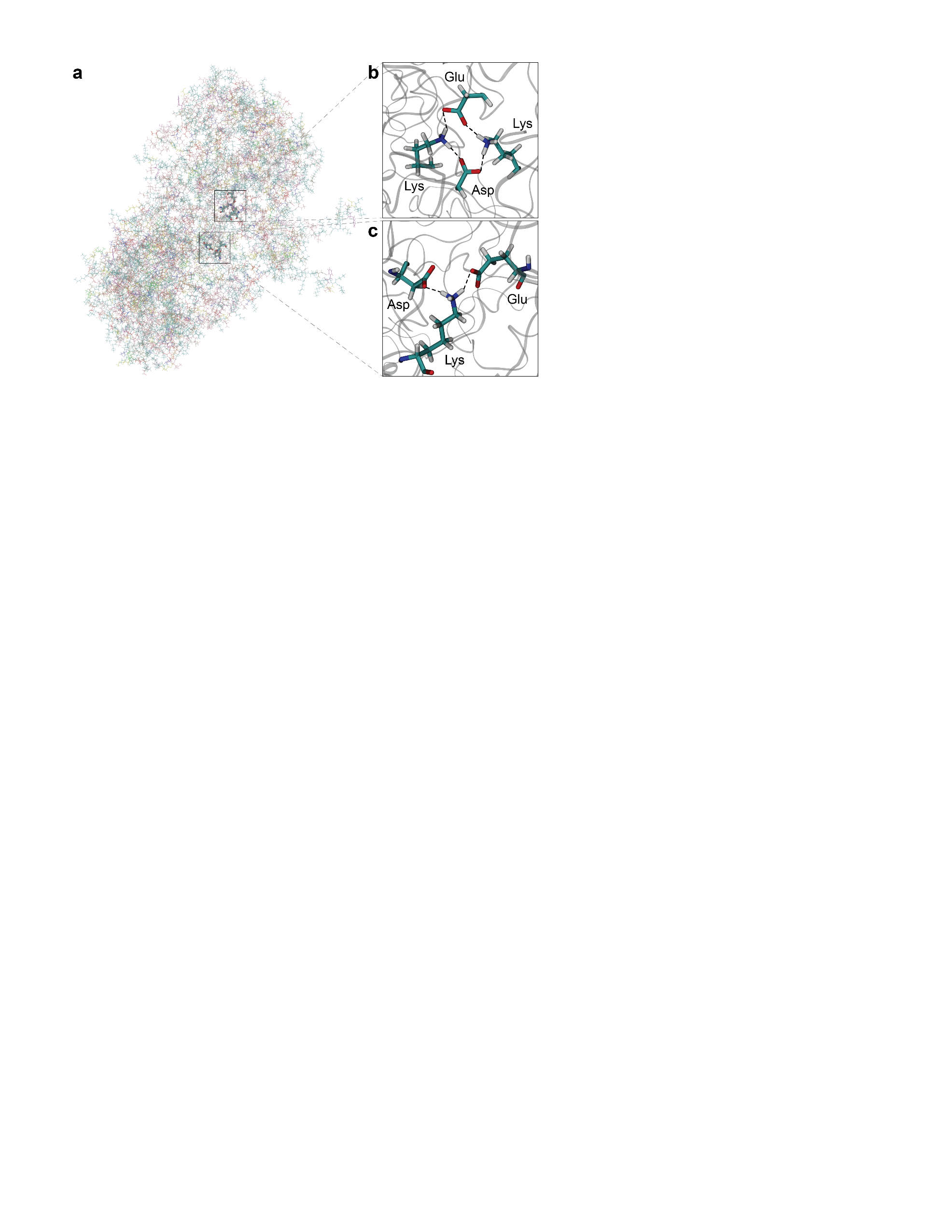

**Supplementary Fig. 4. Atomistic MD simulations predict multiple hydrogen bonds are formed by lysine residue sidechains within amino acid segment 80-95.**

Snapshots from atomistic simulations of the syt1 linker condensate using CHARMM36 force-field^4^. **a** The condensate at atomistic resolution. **b** Magnified view from the interior of the condensate where a complex hydrogen bond (HB) network is shown to involve the sidechains of two Lys, one Asp, and one Glu residue. **c** Another magnified view of the interior of the condensate, where a single Lys sidechain forms simultaneous HBs with Asp and Glu sidechains. These residues are from different copies of the syt1 linker; n = 2 trajectories.

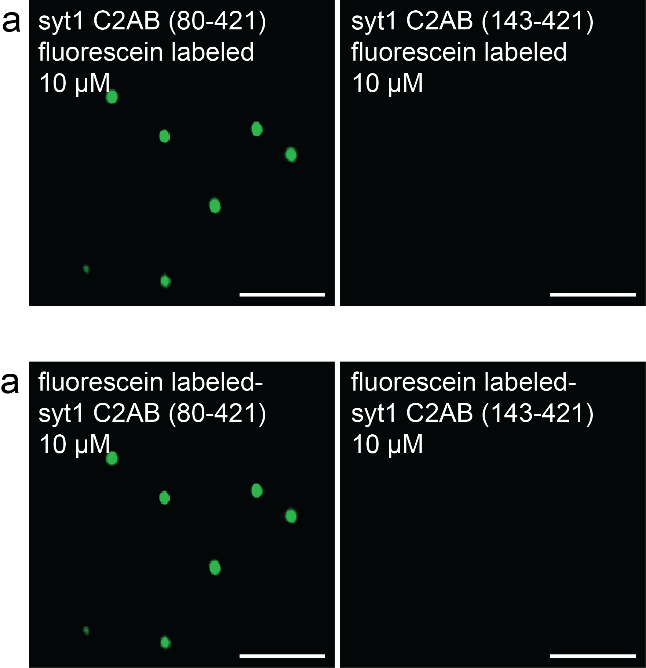

**Supplementary Fig. 5. Fluorescein labeled syt1 forms droplets via the juxtamembrane linker segment.**

**a** (Left) Syt1 C2AB (80-421), labeled with an organic dye (fluorescein) at residues Cys82 and Cys277, formed droplets in the presence of 3% PEG 8000. The buffer condition was 25 mM Tris-HCl (pH 7.4), 100 mM NaCl. The truncated cytoplasmic domain of syt1 (residues 143-421; right), lacking the juxtamembrane linker (residues 80-142) failed to form droplets; n = 3. Scale bars, 3 µm.

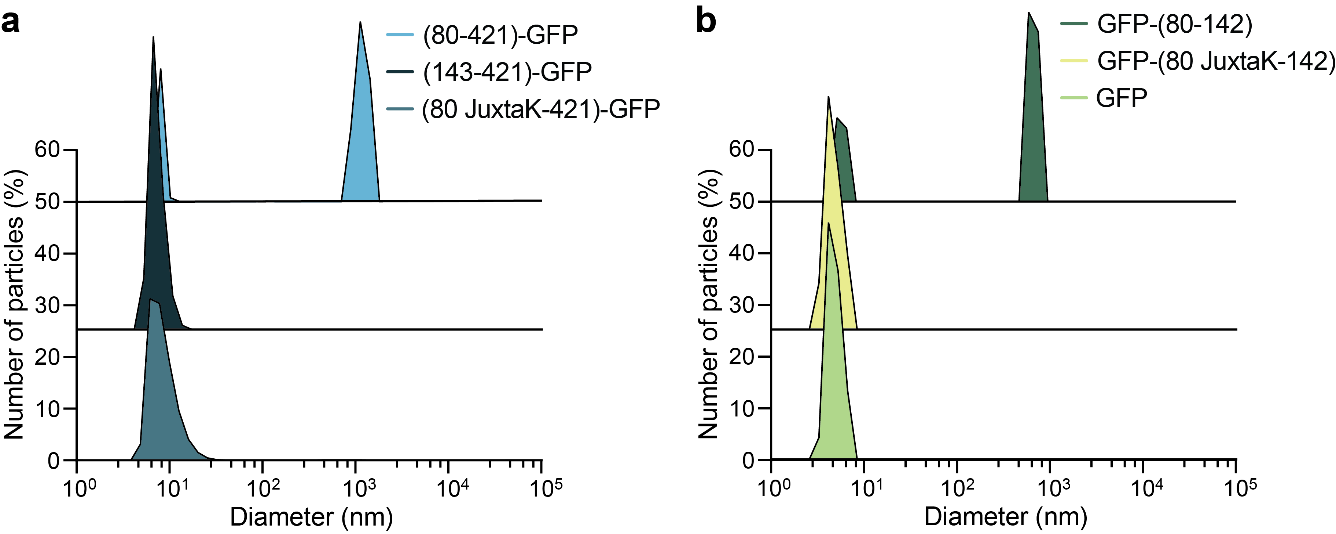

**Supplementary Fig. 6. Syt1 C2AB (80-421)-GFP and GFP-syt1 (80-142) form large oligomers at physiological ionic strength.**

Representative dynamic light scattering (DLS) frequency distributions of the proteins (**Fig. 3a,b). Supplementary Table 3** shows the average diameters; n = 3. The buffer was 25 mM Tris-HCl (pH 7.4), 100 mM NaCl with 3% PEG 8000. Note: these DLS measurements are not accurate at sizes > 1 µm.

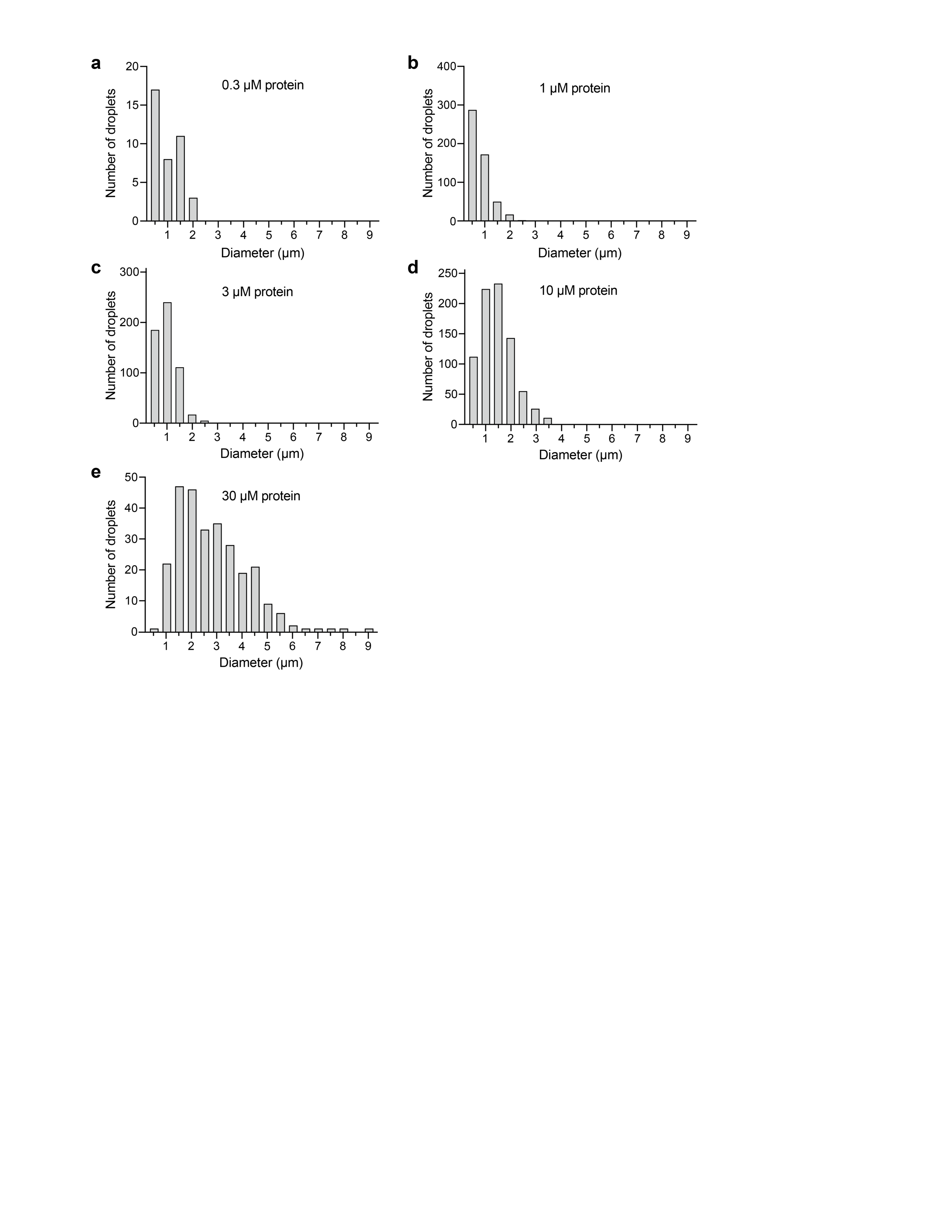

**Supplementary Fig. 7.** **Droplet size correlates with syt1 protein concentration.**

Syt1 C2AB (80-421)-GFP LLPS was monitored at the indicated protein concentrations. At increasing [protein], there is an increase in droplet size. We note that the number of droplets was confounded by the ability of droplets to fuse with each other, so this parameter was not further analyzed. The buffer was 25 mM Tris-HCl (pH 7.4), 100 mM NaCl, 3% PEG 8000; n = 3.

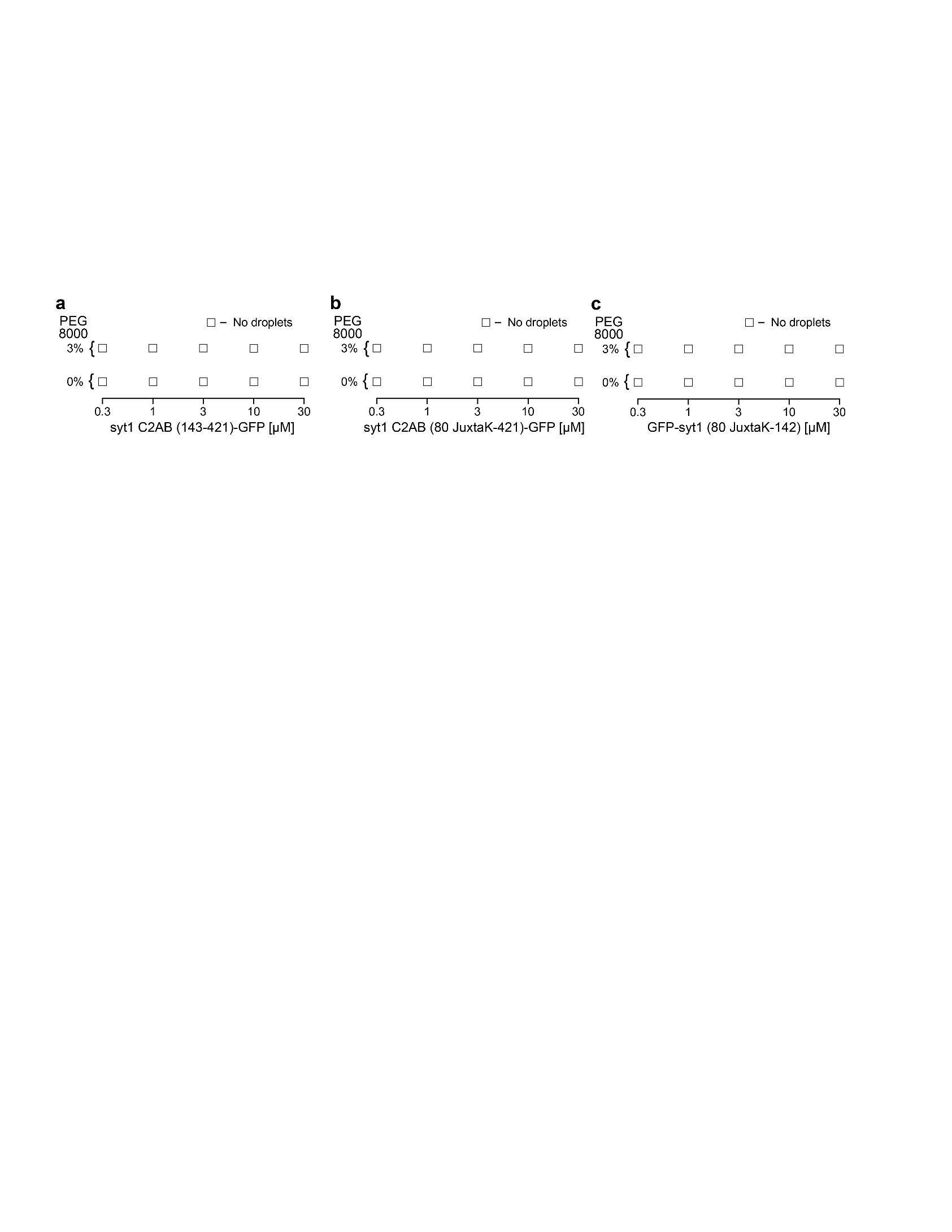

**Supplementary Fig. 8.** **Truncation and mutation of the juxtamembrane linker in syt1 C2AB as well as the isolated JuxtaK mutant linker, fail to form droplets.**

Syt1 C2AB-GFP with a truncated or mutated juxtamembrane linker, as well as the isolated JuxtaK linker, with and without 3% PEG 8000 were monitored for droplet formation, over the indicated [protein] range. White squares indicate that droplets were not observed under any condition tested; n = 3.

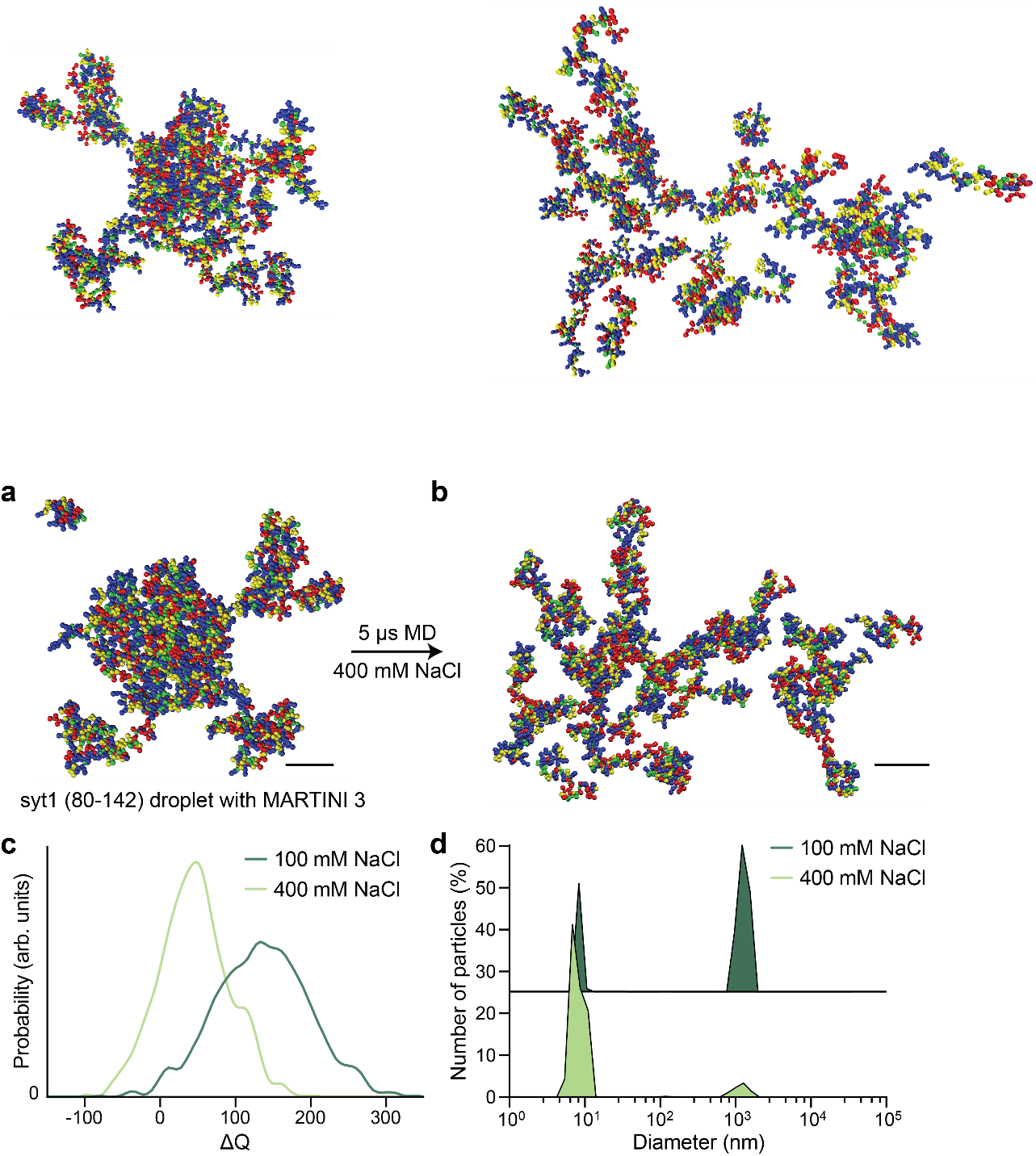

**Supplementary Fig. 9.** **MD simulations predict dissolution of syt1 juxtamembrane linker LLPS droplets at high ionic strength.**

**a** MD simulation of droplet formation by the isolated syt1 (80-142) juxtamembrane linker in 100 mM NaCl after 5 µs of simulation. **b** Droplet partially dissolves at high ionic strength (400 mM NaCl), as determined using the same reparametrized MARTINI v3.0 force field (Scale bars, 4 nm). The color coding is the same as described in Fig. 2c. (See **Supplementary Movie 6** for the movie) **c** The distribution of the gain/loss in the number of contacts (ΔQ) in the syt (80-142) juxtamembrane linker compared to the same number of isolated syt1 linkers. The distribution shifts left at 400 mM [NaCl], indicating a loss in the number of pairwise contacts. **d** DLS analysis reveals the dissolution of syt1 C2AB (80-421)-GFP droplets when the [NaCl] is increased from 100 to 400 mM in 25 mM Tris-HCl (pH 7.4) with 3% PEG 8000. **Supplementary Table 4** shows average diameters and number of particles; n = 3. Source data are provided as a Source Data file.

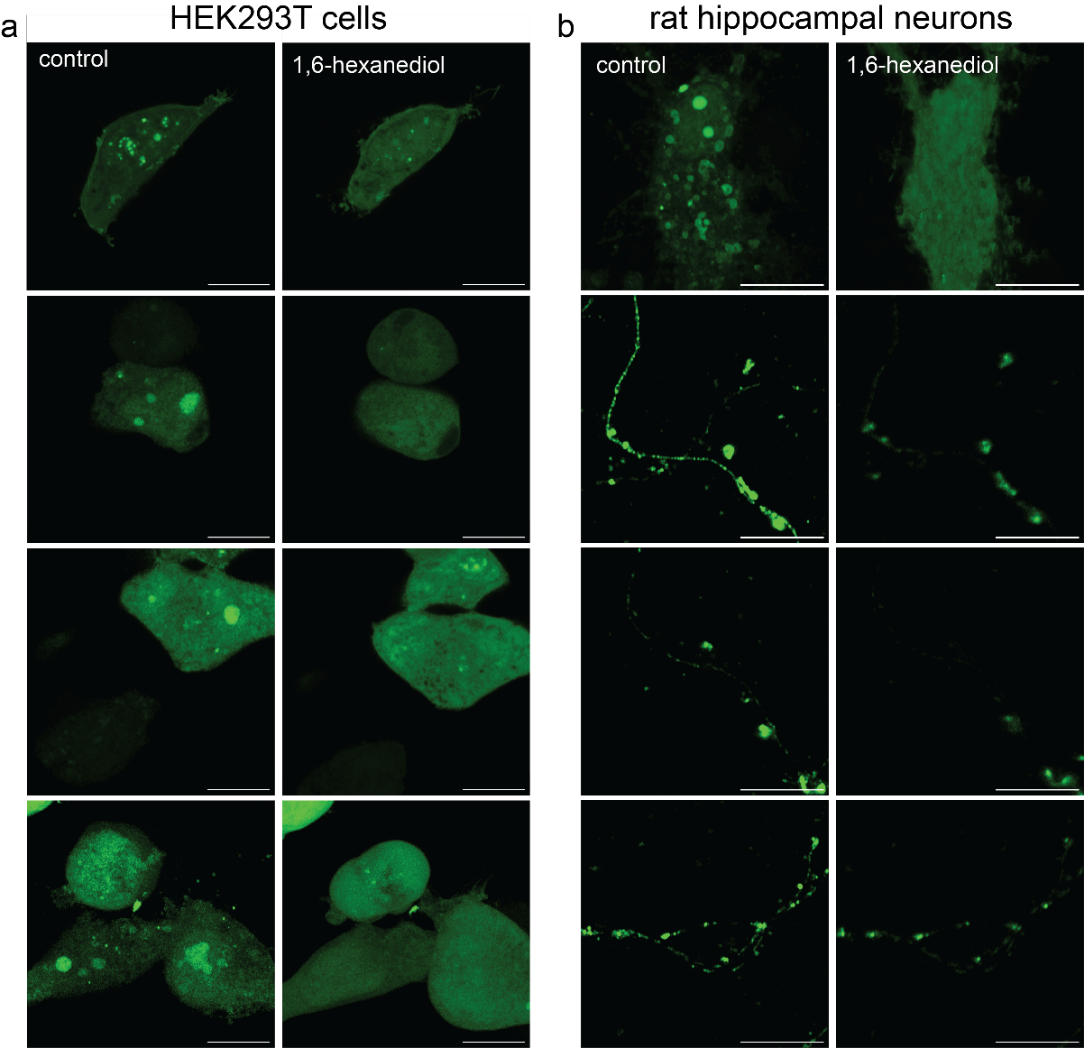

**Supplementary Fig. 10. 1,6-hexanediol dissolves syt1 droplets in HEK293T cells and cultured rat hippocampal neurons.**

**a** Representative super-resolution fluorescence images of HEK293T cells overexpressing syt1 (80-421)-GFP under control and 1,6-hexanediol treatment (5%) conditions. 1,6-hexanediol treatment for 10 min dissolved syt1 droplets. **b** same as **a**, but in cultured rat hippocampal neurons; n = 4. Scale bars, 10 µm.

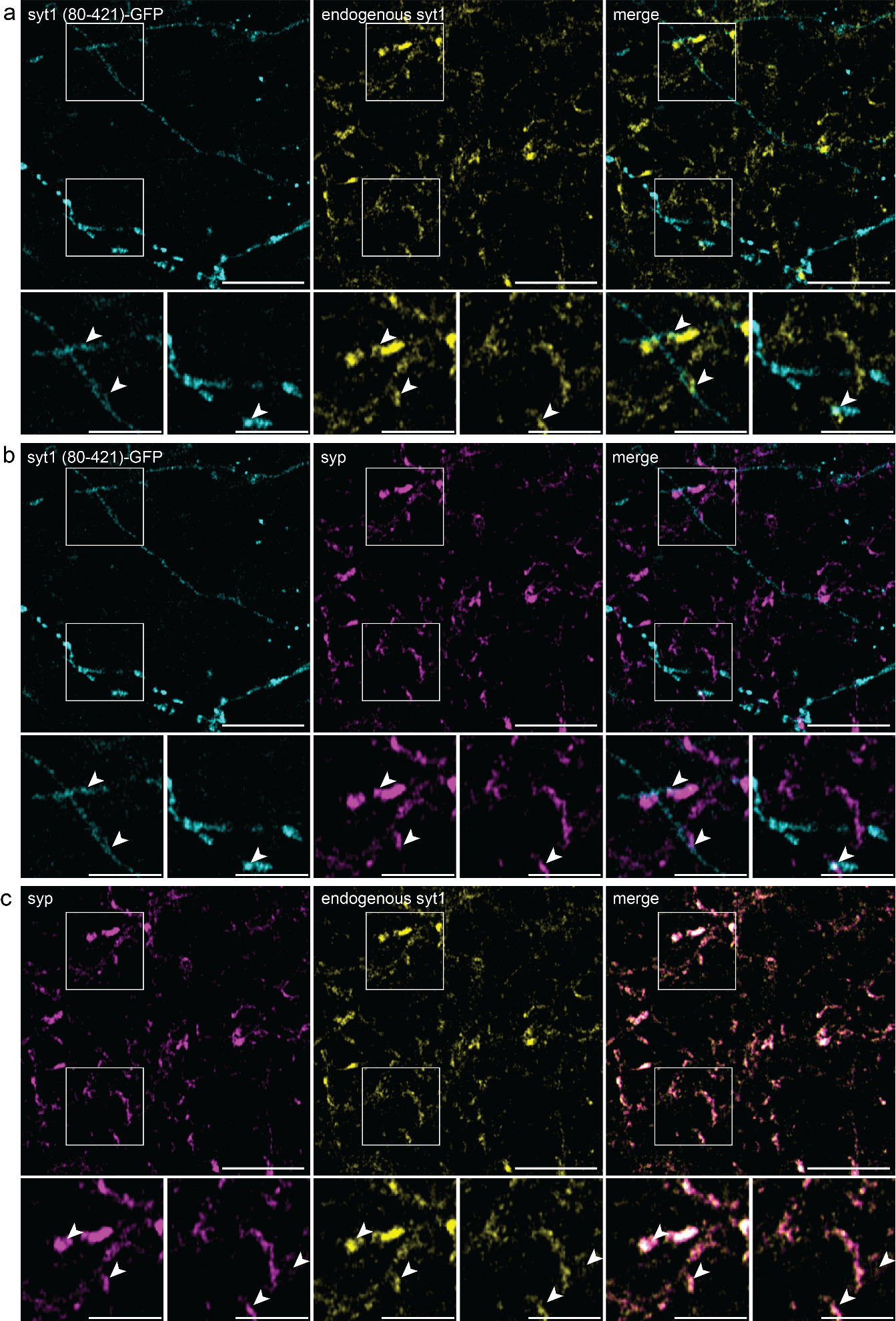

**Supplementary Fig. 11.** **Partial co-localization of syt1 (80-421)-GFP with synaptic vesicle proteins in cultured hippocampal neurons.**

**a** Localization of syt1 (80-421)-GFP was visualized in neurites, whereas endogenous synaptophysin (syp) and syt1 were visualized via immunocytochemistry (ICC); syt1 was selectively detected using an N-terminal luminal domain antibody. Syt1 (80-421)-GFP partially colocalized with endogenous syt1 (**a**) and syp (**b**) as indicated by the arrows in the magnified insets. **c** endogenous syt1 and syp colocalized to a large extent; n = 3. In all cases, images were adjusted with linear brightness and contrast. Scale bars, 10 µm. Inset scale bars, 5 µm.

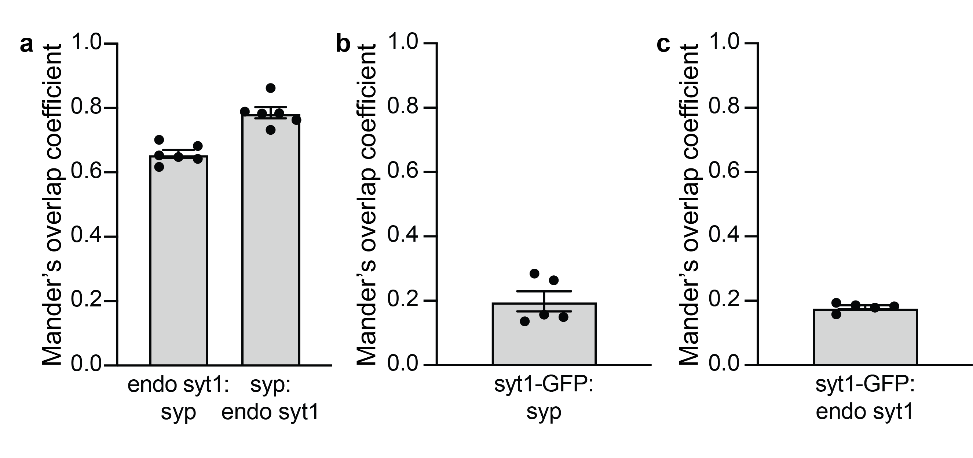

**Supplementary Fig. 12. Colocalization quantification of syt1 (80-421)-GFP, endogenous syt1, and syp**.

**a** Colocalization analysis yields Mander’s overlap coefficient (MOC) of endogenous syt1 and syp as 0.66 ± 0.01 SEM and 0.79 ± 0.02 SEM. **b,c** same as **a**, but for syt1 (80-421)-GFP with endogenous syt1 and syp, yielding MOCs of 0.18 ± 0.01 SEM and 0.20 ± 0.03 SEM respectively. Source data are provided as a Source Data file.

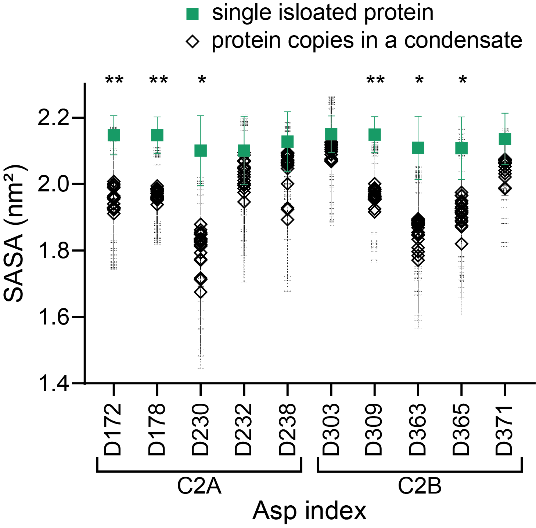

**Supplementary Fig. 13. LLPS masks Ca^2+^-binding to the aspartate residues in the C2-domains of syt1**.

Solvent accessible surface area (SASA) values for the ten aspartate residues proposed to mediate Ca^2+^ binding in the tandem C2-domains of syt1. The green squares denote the SASA values for a single isolated protein and the hollow obliques denote the SASA values for the same aspartate residues in twenty different copies of the protein in a condensate; n = 2 trajectories. Two-sided unpaired t-tests were performed with three datasets using the two-stage step-up method of Benjamini, Krieger, and Yekutieli^5^; p-values of 0.007, 0.006, 0.013, 0.005, 0.012, and 0.03 were obtained for residues D172, D178, D230, D309, D363, and D365 respectively. Source data are provided as a Source Data file.

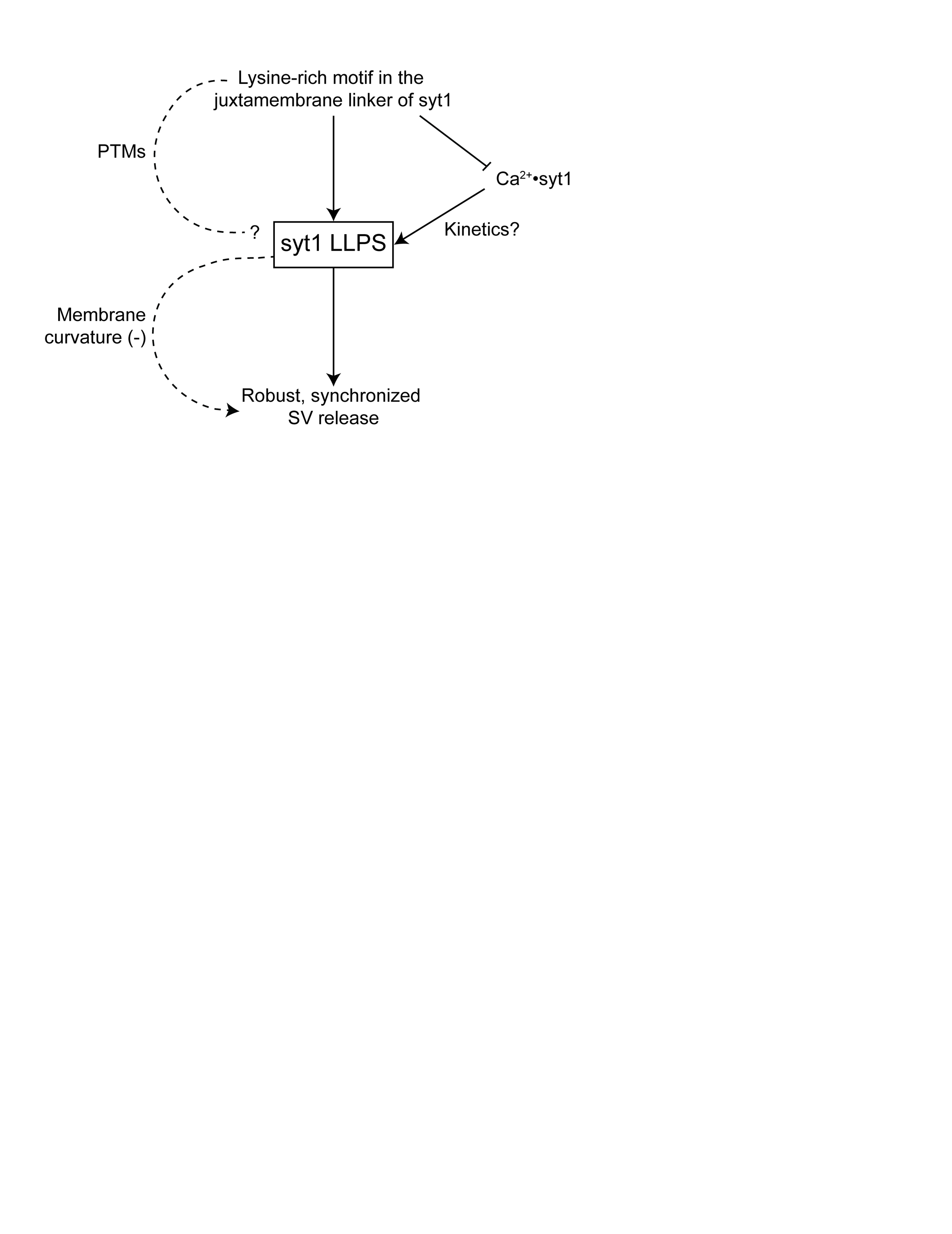

**Supplementary Fig. 14. Model describing how syt1 LLPS might affect syt1 function.**

The lysine-rich motif in the syt1 juxtamembrane linker mediates LLPS (**Fig. 3c,d, 6a,c**), and substitution of these lysine residues disrupts robust, synchronized neurotransmitter release^6^. LLPS reduces the Ca^2+^ sensitivity of syt1, and Ca^2+^ promotes droplet formation. Syt1 LLPS might also favor (-) curvature to facilitate membrane fusion^7^. The juxtamembrane linker harbors sites for post-translational modifications (PTMs) that might regulate LLPS and hence syt1 function. Solid lines represent experimentally established observations; dashed lines indicate proposed phenomenon that are currently under study.

**Supplementary Tables**

|  | DH (cal/mol) | DS (cal/mol/K) | DG (kcal/mol) |
| --- | --- | --- | --- |
| syt1 C2AB (80-421) | DH_1_ = 225 ± 29 | DS_1_ = 19.5 ± 1.2 | DG_1_ = -5.59 ± 0.16 |
|  | DH_2_ = 559 ± 103 | DS_2_ = 18.9 ± 1.3 | DG_2_ = -5.07 ± 0.15 |
|  | DH_3_ = 549 ± 94 | DS_3_ = 17.9 ± 0.3 | DG_3_ = -4.79 ± 0.17 |
|  | DH_4_ = 867 ± 102 | DS_4_ = 16.1 ± 0.7 | DG_4_ = -3.93 ± 0.20 |
|  | DH_5_ = 1040 ± 142 | DS_5_ = 15.1 ± 0.9 | DG_5_ = -3.46 ± 0.23 |
| syt1 C2AB (96-421) | DH_1_ = 96 ± 8 | DS_1_ = 21.5 ± 0.4 | DG_1_ = -6.31 ± 0.02 |
|  | DH_2_ = 618 ± 89 | DS_2_ = 18.9 ± 1.4 | DG_2_ = -5.01 ± 0.14 |
|  | DH_3_ = 661 ± 59 | DS_3_ = 18.7 ± 1.2 | DG_3_ = -4.91 ± 0.09 |
|  | DH_4_ = 832 ± 104 | DS_4_ = 18.1 ± 2.4 | DG_4_ = -4.56 ± 0.03 |
|  | DH_5_ = 1789 ± 102 | DS_5_ = 17.9 ± 2.1 | DG_5_ = -3.55 ± 0.10 |
| syt1 C2AB (80 JuxtaK-421) | DH_1_ = -28 ± 3.1 | DS_1_ = 22.1 ± 1.0 | DG_1_ = -6.61 ± 0.01 |
|  | DH_2_ = -244 ± 51 | DS_2_ = 20.3 ± 0.5 | DG_2_ = -6.29 ± 0.21 |
|  | DH_3_ = -869 ± 256 | DS_3_ = 14.5 ± 1.6 | DG_3_ = -5.19 ± 0.31 |
|  | DH_4_ = -292 ± 101 | DS_4_ = 12.9 ± 1.8 | DG_4_ = -4.14 ± 0.01 |

**Supplementary Table 1**

Thermodynamic properties (G, Gibbs free energy; S, entropy; H, enthalpy) of Ca^2+^ binding to the indicated syt1 cytosolic domain constructs measured using ITC. Data are mean values ± SEM; n = 3.

| **Parameters** | **Expression** | **Value** |
| --- | --- | --- |
| Fraction of positive charges (f_+_) | N_+_/N | 0.301 |
| Fraction of negative charges (f_-_) | N-/N | 0.254 |
| FCR | (f_+_ + f_-_) | 0.555 |
| NCPR | \| f_+_ - f_-_\| | 0.047 |
| SCD | $\frac{1}{N}\left[ \sum_{m=2}^{N} \sum_{n=1}^{m-1} q_{m}q_{n}\left( m-n \right)^{1/2} \right]$ | -21.5 |

**Supplementary Table 2**

Charge composition of the syt1 IDR

Definitions:

a) f_+_ (fraction of positive charges): ratio of the number of positive charges (N_+_) to total number of residues (N).
b) f_-_ (fraction of negative charges): ratio of the number of negative charges (N_-_) to total number of residues (N).
c) FCR (fraction of charged residues): sum of the fraction of positive (f_+_) and negative charges (f_-_).
d) NCPR (net charge per residue): difference between the fraction of positive (f_+_) and negative charges (f_-_).
e) SCD (sequence charge decoration): SCD quantifies charge patterning in a sequence, indicated by the expression in the table. It takes into account the charges (q_m_ and q_n_), along with their separation within the primary protein sequence (m-n).

| Diameter (nm) | syt1 C2AB (80-421)-GFP | syt1 C2AB (143-421)-GFP | syt1 C2AB (80 JuxtaK-421)-GFP | GFP-syt1 (80-142) | GFP-syt1 (80 JuxtaK-142) | GFP |
| --- | --- | --- | --- | --- | --- | --- |
| D1 | 8.8 ± 0.3 (23%) | 8.1 ± 0.6 | 8.2 ± 0.2 | 6.8 ± 0.3 (21%) | 5.8 ± 0.4 | 5.2 ± 0.1 |
| D2 | 1170 ± 91 (67%) |  |  | 891 ± 29 (79%) |  |  |

**Supplementary Table 3**

Table showing the results of the DLS measurements of the syt1 constructs from **Supplementary Fig. 6.** The average diameter of syt1 proteins/oligomers in solution and fraction (%) of small (D1) and large (D2) particles, when two populations were observed; n = 3, error bars represent SEM.

| Diameter (nm) | syt1 C2AB (80-421)-GFP | |
| --- | --- | --- |
|  | 100 mM NaCl | 400 mM NaCl |
| D1 | 8.8 ± 0.3 (23%) | 8.3 ± 0.4 (96%) |
| D2 | 1170 ± 91 (67%) | 1100 ± 75 (4%) |

**Supplementary Table 4**

DLS measurements of syt1 C2AB (80-421)-GFP at low and high ionic strengths from **Supplementary Fig. 9d**. The average diameter of syt1 proteins/oligomers and fraction of particles (%) at 100 and 400 mM NaCl in 25 mM Tris-HCl (pH 7.4) with 3% PEG 8000; n = 3, error bars represent SEM.
